# Organic macromolecules transport a significant proportion of the calcium precursor for nacre formation

**DOI:** 10.1101/2024.01.28.577631

**Authors:** Elena Macías-Sánchez, Xing Huang, Marc G. Willinger, Alejandro Rodríguez-Navarro, Antonio Checa

**Affiliations:** Department of Stratigraphy and Paleontology, Faculty of Sciences, University of Granada, Spain; Department of Chemistry, Fuzhou University, China; Department of Chemistry, TUM School of Natural Sciences, Technical University of Munich, Germany; Department of Mineralogy and Petrology, Faculty of Sciences, University of Granada, Spain; Andalusian Earth Sciences Institute, CSIC−University of Granada, Armilla, Spain

**Keywords:** biomineralization, nacre, surface membrane, vesicles, calcium transport

## Abstract

The mechanism of nacre formation in gastropods involves a vesicular system that transports organic and mineral precursors from the mantle epithelium to the mineralization chamber. Between them lies the surface membrane, a thick organic structure that covers the mineralization chamber and the forming nacre. The surface membrane is a dynamic structure that grows by the addition of vesicles on the outer side and recedes by the formation of interlamellar membranes on the inner side. By using a combination of electron microscopy imaging and spectroscopy, we have monitored the journey of the vesicles from the mantle epithelium to the mineralization chamber, focusing on the elemental composition of the organic structures at each stage. Our data reveal that transport occurs in lipid bilayer vesicles through exocytosis from the outer mantle epithelium. After release into the surface membrane, chitin undergoes a process of self-assembly and interaction with proteins, resulting in progressive changes of the internal structure of the surface membrane until the final structure of the interlamellar membranes is acquired. Finally, these detach from the inner side of the surface membrane. Elemental analysis revealed the transport of a considerable amount of calcium bound to proteins, likely forming calcium-protein complexes.

## 1. Introduction

Nacre is a biomineral that covers the inner surface of the shells of many mollusks, and which shows a characteristic “brick and mortar” arrangement, where the bricks are constituted by calcium carbonate platelets of the polymorph aragonite, and the mortar is constituted by organics. Unlike conventional human-made brick walls, the parallel membranes that separate and provide structure to nacre, also called the *interlamellar membranes* (ILMs), are formed *prior* to the crystallization of the aragonite platelets [1,2]. ILMs are mainly composed of glycoproteins and polysaccharides, specifically chitin in β-sheet conformation [3-5]. Constituting only 5% of the shell dry weight, this organic component confers the biomineral a precisely controlled hierarchical organization, that results in a resistance to fracture significantly superior to its inorganic counterpart [6].

Two out the four nacre-forming groups, bivalves and gastropods, have evolved different nacre production mechanisms. In the case of bivalves, nacre precursors are secreted by the ectodermic cells of the mantle epithelium to the *extrapallial space*, a narrow cavity (∼50-100 nm) between the tips of the mantle cell microvilli and the shell growth surface. This enclosed space provides the controlled environment necessary for the self-assembly of the shell components.

However, gastropods do not have a stationary extrapallial space, but it is bounded by the soft body, and therefore subject to the mobility of the organism. The formation of the ILMs derives from a dense superficial layer, called the *surface membrane*, which lines the external part of the mineralization compartment [1]. The side of the surface membrane adjacent to the mantle epithelium is composed of a dense organic material, formed by the addition of vesicles [7]. However, in the innermost region, facing the mineralization compartment, the structure changes towards an arrangement of seemingly welded membranes [1,8] from which fresh ILMs progressively detach in a zipper-like fashion [2,7] and are positioned with a typical spacing of about 500 nm [9].

In recent decades, attention has focused on the transport and transformation of amorphous phases for the formation of biominerals. The case of nacre is no different, and the presence of amorphous calcium carbonate (ACC) has been demonstrated in early stages of mineralization of gastropod nacre [10]. However, little attention has been devoted to the structure-function relationships of the organic structures (interlamellar membranes, surface membranes) that precede mineralization. Using electron microscopy and spectroscopy we have studied the early stages of mineralization in three marine gastropods, *Phorcus turbinatus, Steromphala pennanti* and *Steromphala* sp. (Trochidae, Vetigastropoda).

We have monitored the vesicular transport system, from the mantle epithelium to the self-assembly of the polysaccharide/protein scaffold at the surface membrane inner side. Our data demonstrate that a significant amount of calcium is transported bound to organic macromolecules in the form of calcium-protein complexes, which then associate with β-chitin to provide the organic framework for aragonite platelet mineralization.

## 2. Methods

### 2.1 Critical point drying for SEM observations

Specimens of *Steromphala pennanti* and *Steromphala* sp. were caught alive in Almuñécar (36º44’02’’N, 3º41’28’’O), in the coast of Granada province (Spain), fixed alive (glutaraldehyde 2.5% in cacodylate buffer pH 7.4) and prepared by critical point drying. SEM investigation was carried out in a Zeiss Gemini at the Scientific Instrumentation Center of the University of Granada (CIC-UGR).

### 2.2 Chemical fixation

Specimens of the nacreous gastropod *Phorcus turbinatus* were collected in Benalmádena (Málaga, Spain 36°35’42’’N, 4°30’56’’W). Immediately after capture, specimens were fixed in cacodylate-buffered glutaraldehyde 2.5%. Fragments of the surface membrane area were fractured under the binocular microscope, postfixed in 1% osmium tetroxide, dehydrated in ethanol series and embedded in epoxy (EMbed 812, Electron Microscopy Sciences).

### 2.3 Histology of the mantle

*Phorcus turbinatus* specimens were caught alive at the same spot, transported to the lab, anesthetized by intradermal injection of MgCl_2_ and dissected with a scalpel to isolate the mantle. Mantle tissue was sectioned into 1-2 mm pieces and fixed in cacodylate-buffered 2% paraformaldehyde and 4% glutaraldehyde, postfixed in 1% osmium tetroxide, dehydrated in ethanol series and embedded in epoxy.

### 2.4 Flash-freeze and freeze-drying

Juvenile specimens of *Phorcus turbinatus* were caught alive in La Herradura (36°43’44’’N, 3°43’35’’W), in the coast of Granada province (SE Spain), frozen *in situ* in liquid nitrogen and stored in a vacuum flask until the next day. Freeze-drying was carried out in a Flexi-Dry MP (CIC-UGR). The initial temperature (−170°C) was raised slowly at an average pressure of 75 mTorr for 2 days. At the end of the drying cycle (20°C), the specimens were removed and stored at 4°C. This process involves exclusively the elimination of unbound water (sublimation phase or primary drying).

Specimens were cut with a diamond saw and pieces from the surface membrane area were selected for embedding with epoxy (Embed 812, EMS), increasing the proportion of embedding medium to pure ethanol in three steps (1:2, 1:1, and 2:1). Osmication was applied to half of the samples and avoided for the rest. Ultrathin sections were cut (PowerTome Ultramicrotome) at a small angle relative to the surface (< 10º), thus maximizing the visualization likelihood of the surface membrane and interlamellar membranes. The slices were laid in copper grids with lacey carbon to stabilize them under the electron beam.

### 2.5 Transmission Electron Microscopy (TEM) Imaging

Transmission Electron Microscopy imaging was carried out at 200 kV using either a Philips CM200 equipped with a field emission gun and a Gatan UltraScan 4000 GIF camera or a double Cs-corrected JEOL JEM-ARM200F equipped with a Gatan Oneview camera (Fritz Haber Institute of the Max Planck Society, Berlin).

### 2.6 Energy Dispersive X-Ray Spectroscopy (EDS)

EDS mapping was acquired at 200 kV using a beryllium holder on a Cs-corrected FEI Titan (ThermoFisher) equipped with a field emission gun and 4 silicon drift detectors for EDS analysis (Super-X detection system) (CIC-UGR). Spectra were exported to Digital Micrograph for further analysis using the script “Read Brucker EDS spectrum text file” of Dave Mitchell.

### 2.7 Electron Energy Loss Spectroscopy (EELS)

EEL spectra were obtained at 200 kV by means of a post-column energy filter spectrometer (Ultrascan 4000 GIF camera, Gatan) using a beryllium double tilt holder. Emission current was set to 5 µA to increase energy resolution and reduce the total dose. EEL spectra covering C Kα, Ca L_2_-L_3_, N Kα and O Kα core edges (from 240 to 650 eV) were recorded using a dispersion of 0.2 eV/channel, with a collection semi-angle β = 20 mrad and a 2 mm spectrometer entrance aperture. The energy resolution was ∼1.4 eV measured by the full width at the half maximum (FWHM) in the zero-loss peak.

For spectra covering one edge (Ca L_2-3_), we used a dispersion of 0.05 eV/channel, with a collection semi-angle of β = 20 mrad and a 2 mm aperture. The energy resolution was ∼0.95 eV at FWHM in the zero-loss peak.

Sample thickness was computed using the relative log-ratio method [11] and in all cases t/λ < 0.2 (between 0.14 and 0.2), and therefore plural scattering contribution can be considered negligible. In any case, the background was subtracted using power-law fitting before each edge and plural scattering was removed using a Fourier-ratio deconvolution, both available in Digital Micrograph 3.1 software (Gatan Inc.). To reduce the noise, addition of 5 spectrum plus 2-point smoothing was used in Figure 5. Data plotting and normalization were performed using Origin Pro 9.

### 2.8 Fourier Transform Infrared Spectroscopy (FTIR)

Fragments of the nacre layer of *Haliotis rufescens* and *Phorcus turbinatus* (in the latter case containing the surface membrane) were demineralized in 4% EDTA. Demineralized interlamellar membranes were carefully located on a glass slide and air dried. FTIR spectra were recorded with a JASCO 6200 instrument (CIC-UGR) fitted with an ATR device in the 4000-600 cm^-1^ wavenumber range at 0.5 cm^-1^ spectral resolution.

## 3. Results

### 3.1 The system

The growth of the nacreous layer takes place near the shell aperture [12,13]. It is preceded by an aragonitic fibrous prismatic layer at the very shell edge and followed (and covered internally) by a platy aragonitic material with nacreous luster (Figure 1a-b, Supplementary Figure S1). In transversal cross-section, the surface membrane appears as a dense organic structure covering the growing nacre (Figure 1c, d). Interestingly, the tips of the incipient towers are always in contact with the surface membrane (Figure 1d) from where the growing tablet absorbs organic components [7]. On the mantle side, the surface membrane appears covered by a multitude of vesicles that gradually integrate into the whole structure, whereas on the nacre side, the interlamellar membranes detach from it (Figure 1e). The spaces delimited by the interlamellar membranes where the crystals grow are filled by a silk-like protein with a gel-like structure containing acidic glycoproteins [14,15]. When prepared by critical point drying, this organic filler material dehydrates and aggregates, appearing dispersed on the interlamellar membranes (Figure e-h).

**Figure 1.**
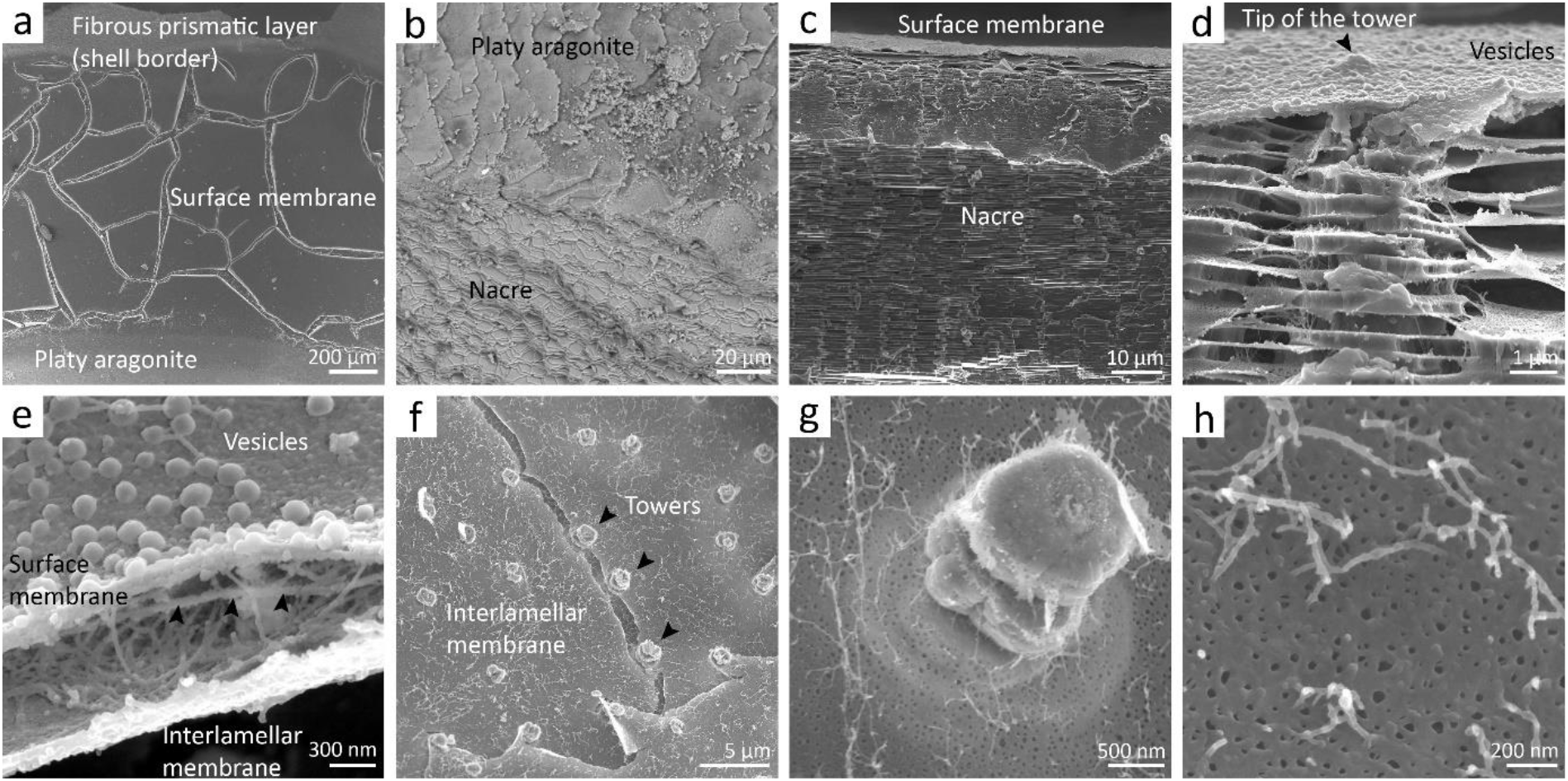
General view of the system. a) A fibrous prismatic layer grows on the edge of the shell, followed to the shell interior by the surface membrane, with a smooth and homogeneous appearance (the cracks are artifacts of sample preparation). A layer of platy aragonite grows next. b) Detail of the boundary between the platy aragonite and the underlying nacre. c) Cross section of the growing nacre. The material is growing actively in the upper zone, where the platelets are incomplete. d) The tips of the growing towers are always immersed in the surface membrane. Each platelet grows within the space limited by the interlamellar membranes. e) Surface membrane covered with vesicles. Vertical fracture shows the last two platelets formed and two interlamellar membranes, the one closest to the surface membrane in the process of separation (arrowheads). A large amount of organic matter is observed between the membranes. f) Top view of the mineralization chamber with the surface membrane removed showing the two most recent interlamellar membranes and nacre towers. After critical point drying, the organic matter that fills the spaces between the interlamellar membranes in the native state, precipitates on the membranes. g) Detail of a tower and an intact interlamellar membrane. h) Detail of the organic material precipitated on the interlamellar membranes.

### 3.2 Vesicle system

The mantle of gastropods consists of two folds (the inner mantle fold and the outer mantle fold) separated by the periostracal groove (Figure 2a). Within the periostracal groove we found an additional accessory fold (Figure 2a-b), not found in other species. The inner mantle fold (Figure 2c) is characterized by the alternation of secretory and non-secretory cells and by conspicuous cell-adhesive unions (Figure 2c, arrowheads). The outer shell-facing epithelium (also known as the *outer mantle epithelium*) is the zone responsible for shell formation [16]. It consists of a single layer of epidermal cells with short microvilli forming a folded epithelium with an undulating surface (Figure 2d) and is constituted by three main types of cells: microvillous epidermal cells, ciliated cells and secretory cells [17,18]. The mantle ectodermal cells form an extremely active secretory epithelium, extruding an extraordinary number of vesicles (Figure 2e). The great diversity of vesicle sizes and shapes suggests a complex system controlling the secretion of organic and mineral precursors.

**Figure 2.**
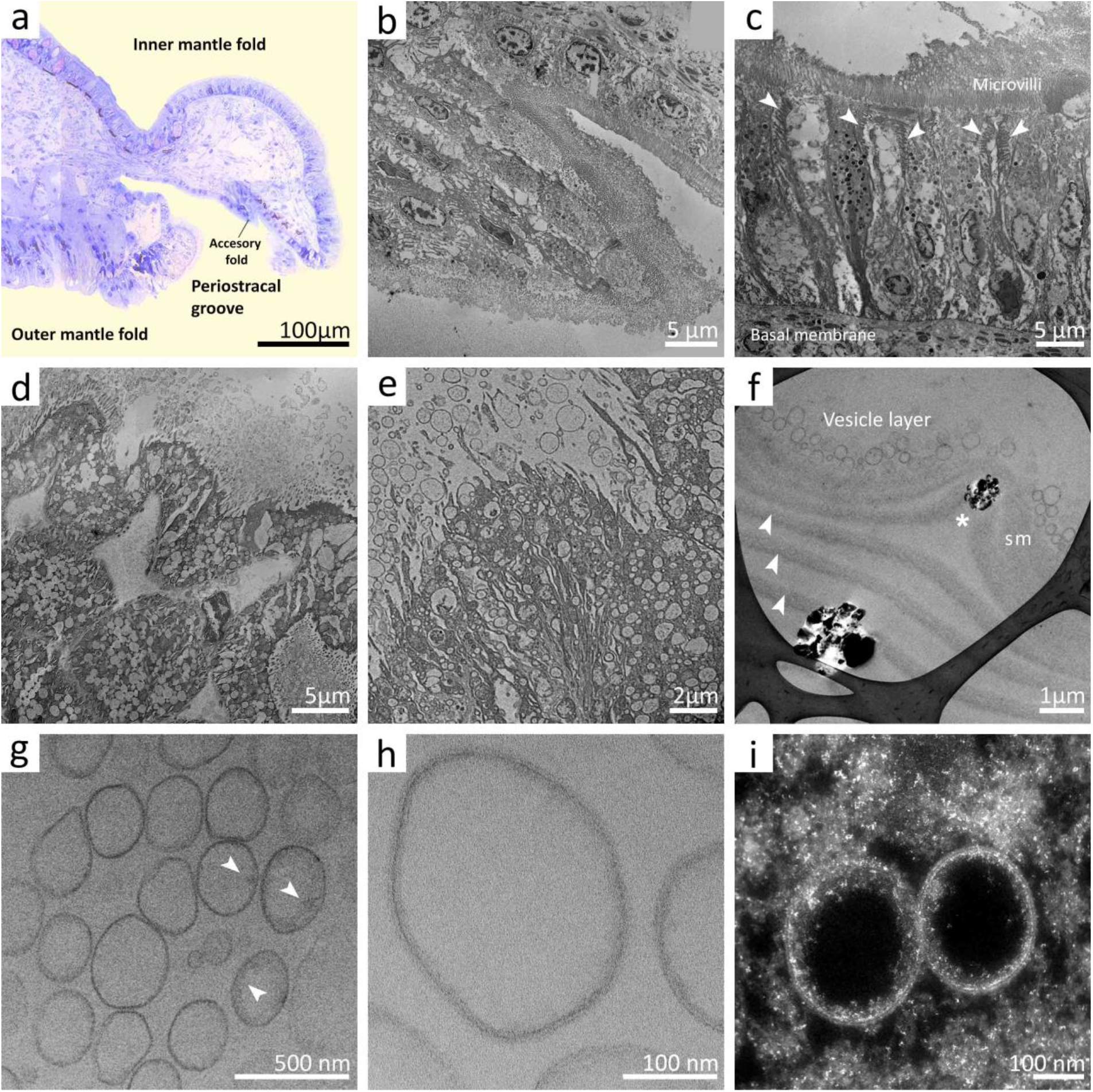
Vesicle system: from the mantle epithelium to the surface membrane. a) Histological slide of resin-embedded mantle epithelium stained with toluidine blue. The inner mantle fold, the outer mantle fold, the periostracal groove, and the accessory fold are indicated. b) Detail of the accessory fold located in the periostracal groove. c) Image stitching showing the monolayer of cells with dense microvilli in contact with the basal membrane characteristic of the inner mantle fold. Secretory and non-secretory cells alternate. d) Folded epithelium characteristic of the outer mantle fold. e) Microvilli of the cell apical area showing the characteristic active exocytosis process. f) On the side of the mineralization compartment, the tips of the towers (marked with an asterisk) appear immersed in the surface membrane (sm) and a large number of vesicles. The interlamellar membranes (arrowheads) detach from it. g) Cargo attached to the inner membrane of the vesicles (arrowheads). h) Detail of the lipid bilayer (bright field TEM image). i) HAADF-STEM provides high contrast images based on atomic number differences (Z-contrast). In this case, the osmium tetroxide attached to the lipids double bounds and organic material produces a high contrast against the carbon-based epoxy resin, revealing the bilayer structure of the membrane. When the vesicles release the transported material, it aggregates surrounding the remaining vesicles.

A layer of vesicles invariably appeared on the external side of the surface membrane (Figures 1e, 2f). They were mostly round with an average size of ∼200 nm (ranging from 50 to 400 nm) and showed the characteristic bilipid membrane (10-15 nm thick) (Figure 2g-i). The cargo was found to be preferentially bound to the inner membrane (Figure 2f, h-i; see also Figure 3b-c). After fusion of the vesicles with the surface membrane, the transported materials diffuse, forming an accumulation of amorphous organic material that mixes with the remaining vesicles (Figure 2i).

**Figure 3.**
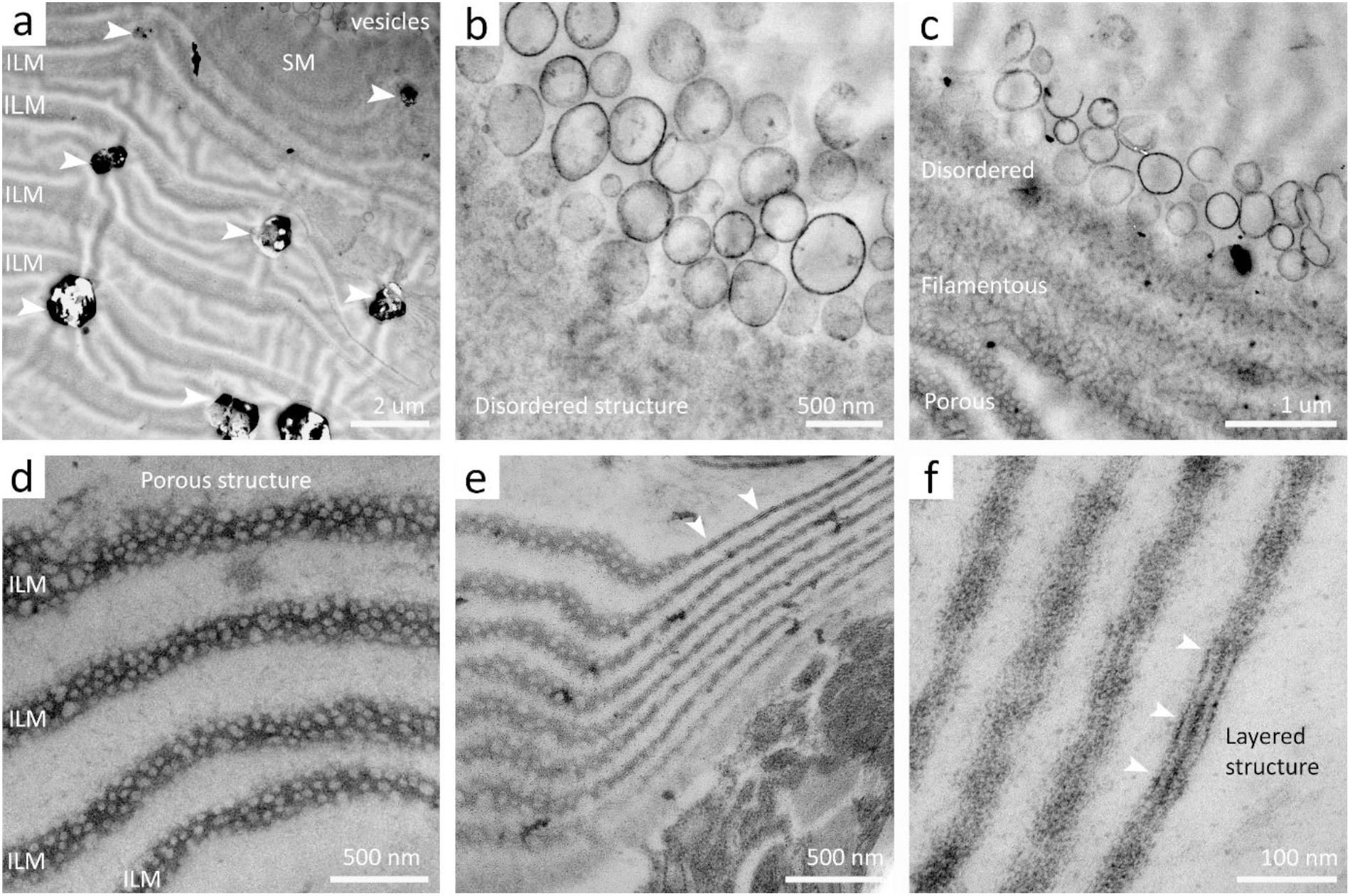
Polymerization of the surface membrane. a) Overview of the system showing the sequential organization from the vesicle layer and the surface membrane (SM) to the interlamellar membranes (ILM). Incipient platelets are indicated by arrowheads. b) Vesicle fusion and cargo release generates a zone of disordered structure. c) Transition from a disordered into a filamentous and, ultimately, a porous structure. d) Porous structure characteristic of gastropod interlamellar membranes (diagonal section). e) Twisting of the interlamellar membranes allows their different projections to be observed in the same image: on the left, the characteristic porous structure visible in diagonal section; on the right, the characteristic three-layer structure observed in transversal cross-section. f) Detail of the three-layer structure. a-c) Inverse HAADF STEM; d-f) BF TEM, post-stained with uranyl acetate and lead citrate.

### 3.3 Surface membrane polymerization

Vesicles extruded from the mantle epithelium reach the surface membrane that covers the mineralization compartment. Underneath, the nacre platelets grow in towers within the spaces defined by the interlamellar membranes (Figure 1d; Figure 3a, diagonal section). The formation of the surface membrane results from the fusion of a multitude of vesicles and the release of their cargo (Figure 3b-c). When observed in cross-section, the change in the structure of the surface membrane appears to be the result of a process of polymerization of the transported precursors. Three structurally distinct regions were identified across (indicated in Figure 3c). (1) The outermost region, facing the mantle and next to the vesicular layer, which showed a *disordered* and diffuse structure (Figure 3b). (2) Towards the mineralization compartment, we found an intermediate region characterized by a *filamentous structure* (Figure 3c). In the innermost region, the surface membrane acquired a *porous structure*, similar to that of the interlamellar membranes (Figure 3c, d). This is also the zone of detachment of the fresh ILMs.

It is important to note that the porous structure is evident in plan-view and in diagonal cross-section (Figure 3d). When cut in transversal cross-section, the interlamellar membranes show the characteristic three-layered structure (Figure 2e, f), as reported previously [1,19].

### 3.4 Calcium transport

Only flash-frozen/freeze-dried samples retained the calcium content, the signal intensity being higher when OsO_4_ was used. Samples prepared by traditional chemical fixation retrieved no calcium signal, probably due to the washout of ionic species and mineral precursors during the different aqueous steps of sample preparation [20-22].

We conducted a pre-screening using STEM-EDS to determine the spatial distribution of calcium in the organic osmium-stained system. EDS mapping revealed the presence of calcium in the vesicles (membrane and cargo), the surface membrane and downstream ILMs (Figure 4a-b). These areas were subsequently used to perform systematic EELS measurements.

**Figure 4.**
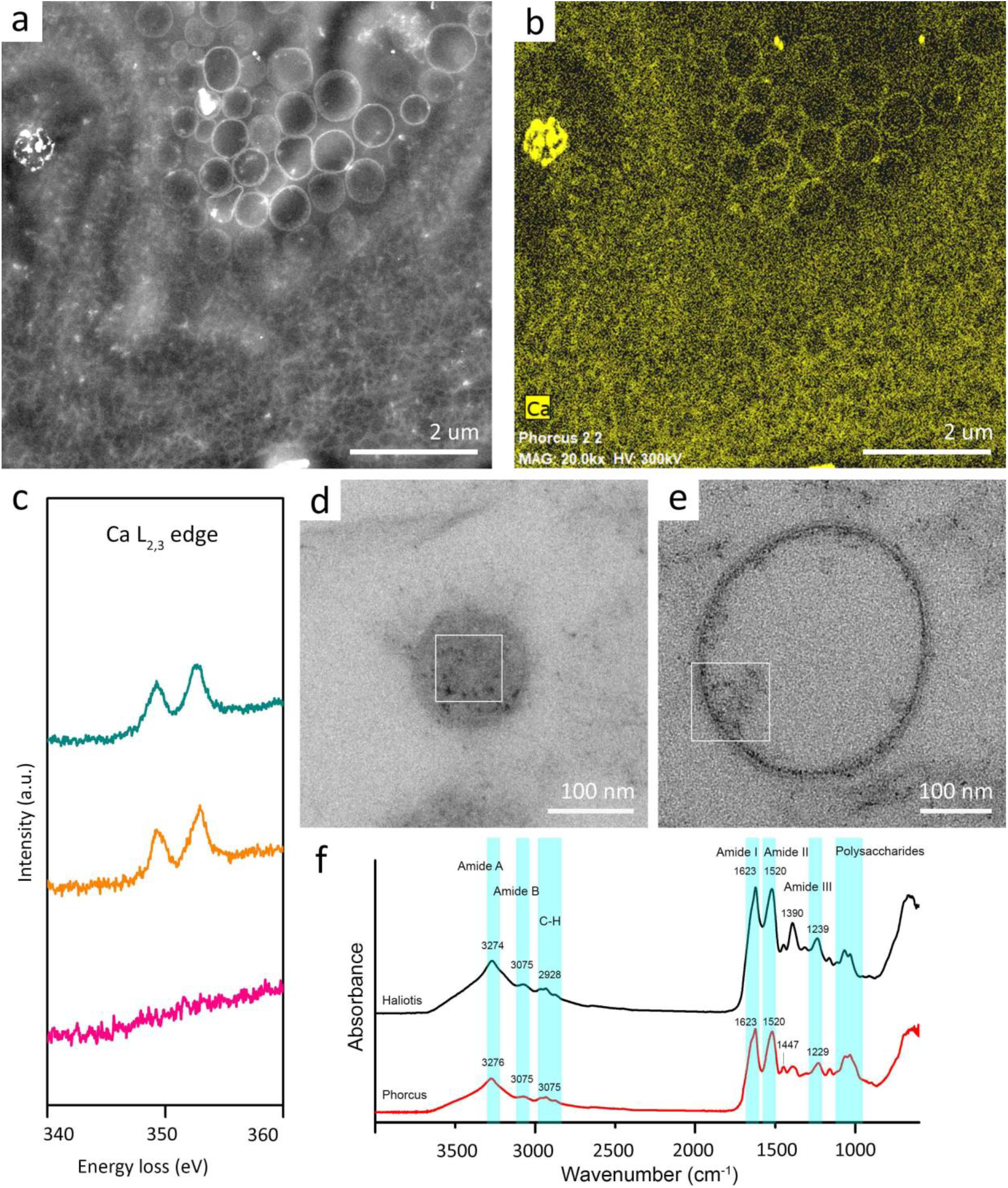
The organic content of the vesicles incorporates a certain amount of calcium. a) HAADF-STEM image of the vesicle layer, the surface membrane and the interlamellar membranes. b) EDS-STEM elemental mapping from the same area showing the calcium distribution in the system. c) Comparative Ca L_3,2_-edge of three different areas: vesicle cargo (green), vesicle membrane (orange) and resin (pink). d-e) TEM images showing different examples of measured vesicles: d) a completely filled vesicle and e) partially filled vesicles, where the material appears adhered to the inner side of the membrane. The squares indicate the measured area. f) FTIR spectra from the demineralized nacre layer of *Haliotis rufescens* (black) and *Phorcus turbinatus* (red).

The epoxy-only regions showed a C K-edge constituted by an initial peak at 285 eV and a second broad feature at 290 eV, characteristic of amorphous carbon. Small features at the N K-edge (400 eV) and O K-edge (532 eV) indicate a minor amount of these elements, as expected from the resin chemical composition (Supplementary Figure S2). No calcium signal was measured. Considering that a significant contribution to the carbon peak comes from the resin, we avoided further analysis of this edge. Vesicle cargo retrieved a significant calcium signal (Ca L_2,3_ edge, 349.3 eV), while the vesicle membranes also reported a calcium signal, although of lower intensity (Figure 4c).

Comparison of the FTIR spectra from *Haliotis rufescens* and *Phorcus turbinatus*, the latter including the surface membrane, showed an almost identical profile. The amide I signal retrieved from both samples consists of a single band at 1623 cm^-1^, characteristic of β-chitin [23,24]. The location and relative intensities of the rest of the amide bands (amide II, III, A and B) were similar (1520 cm^-1^, 1239/1229 cm^-1^, 3274/3276 cm^-1^, 3075 cm^-1^), as well as the peaks in the range 950-1110 cm^-1^ related concentration of glycosidic bonds, and the ν(O-H) stretching vibration mode of the hydroxyl OH groups. The only appreciable differences between both spectra relates with the relative intensities of the C-H scissoring vibration mode (1447 and 1390 cm^-1^). Taken together, the data indicate that the composition of the surface membrane (present in the *Phorcus turbinatus* sample) does not differ chemically from that of the interlamellar membranes.

The interlamellar membranes showed an intense calcium signal (Ca L_2,3_ edge, ∼349.3 eV) and a significant amount of nitrogen (N K-edge, 400 eV), hallmark of proteins. Interestingly, the electrodense organic aggregates dispersed between the interlamellar membranes also showed both calcium and nitrogen signals, although with lower intensity.

## 4. Discussion

### 4.1 Vesicle system

The thickness of the vesicle double membrane and the intense OsO_4_ staining (Figure 2e-i), which binds preferentially to double bounds of phospholipids, indicate that they are endosomes constituted of a lipidic bilayer and formed by strangulation of the cell membrane (Figure 2d-e).

The content of the vesicles was variable in amount, ranging from full to partially empty vesicles, and was mostly attached to the inner membrane (Figure 2e-i; Figure 3b-c; Figure 4a, d-g). Although we could identify calcium within the vesicles, not all of them yielded calcium signal, supporting previous proposals that different precursors may be secreted independently by specialized mantle cells restricted to discrete areas [16].

### 4.2 Surface membrane: chitin self-assembly and formation of the ILMs

Far from being a static structure, the surface membrane is a dynamic entity, assembled by the addition of vesicles and their cargo at the mantle side, and deconstructed at the mineralization chamber side, by the separation of the interlamellar membranes (Figure 6).

It is well established that the interlamellar membranes are mainly composed of proteins and β-chitin [3], with a small amount of lipids (0.54% by weight) [25], which, in the case of gastropods, must derive from the vesicles involved in the transport system. We expected the composition of the surface membrane to be similar, as it constitutes a preliminary stage of the interlamellar membranes, what has been supported by FTIR data. In fact, the surface membrane represents the *medium* where the chitin self-assembly and the interaction with the proteins takes place. The optimal pH range for the formation of chitin microfibers ranges from 7 to 8.5, which includes not only the pH of physiological conditions but also that of seawater [26]. The pH of the extrapallial fluid in marine bivalve species ranges from 7.2 to 7.4 [27] and up to 8.3 in freshwater species [28]. It is important to note that in gastropods there is not a fixed extrapallial space, and in response to external stimuli, the animal retracts inside the shell leaving the growing nacre compartment exposed to seawater (Supplementary Figure S3). This behavior does not seem to alter the formation process, so the surface membrane should be impermeable enough to keep the necessary concentrations of organic and mineral precursors for the formation of the interlamellar membranes and the subsequent mineralization of nacre platelets. SEM images showed the outer surface of the surface membrane as a fibrous structure embedded in a dense protein matrix, with no apparent pores (Figure 1d-e), although we cannot rule out if this appearance is an effect of the critical point drying.

Moving in depth, the structure changes acquiring the characteristic fibrous and porous structure, probably driven by a reorganization of the protein layer (Figure 3c, Figure 6). Chitin-protein interaction occurs through hydrogen bonds, once chitin polymerization has concluded [29] and appears to protect the nanofibrils against disassembly and biodegradation [26]. This chitin-protein network is in turn embedded in an additional protein matrix [30].

It should be noted that the tips of the towers are always embedded within the surface membrane [7]. In addition, previous work has highlighted that the nuclei of gastropods tablets are enriched with organic material [13,31] and constitute a continuous central axis across superposed tablets [7]. Consequently, the organic material that composes the central core of the plates should be absorbed from the surface membrane. In other groups (*Atrina*, Bivalvia; *Nautilus*, Cephalopoda) the core mainly consists of acidic proteins [32] that appear to be concentrically distributed [33].

### 4.3 Calcium transport

The osmium-stained organic material retrieved a clear calcium signal throughout the entire transport system, indicating that a substantial proportion of the calcium was bound to organic macromolecules. Osmium tetroxide preferentially binds to the C=C double bonds, i.e. phospholipids, but also to some proteins containing certain amino acids (tryptophane, cysteine, histidine) and unoxidized sulfur (e.g. sulfhydryl groups) [21,34], which may relate with the presence of sulfated glycoproteins. Furthermore, the C K edge of the EELS spectra did not yield the characteristic carbonate peak at 290 eV, but only amorphous carbon (Supplementary Figure S2), indicating that the calcium content bound to organic matter is in organic form.

Of particular interest is the presence of calcium in the organic threads scattered between the interlamellar membranes (Figure 1e-h; Figure 5c). Nakahara [1] first noted the presence of organic aggregates between the interlamellar membranes, indicating that they form a network associated with the organic sheets covering the sides of the growing crystals. According to their nitrogen signature, our results show that they are proteins extruded by the mantle and transported by vesicles. These results are in line to previous reports which indicated that 74.3% of the calcium in the extrapallial space of *Mytilus edulis* was bound to small chelates, and 9.2% was tightly bound to insoluble sulphated carbohydrates [27]. Our data are in line with recent reports that showed the existence of an outer amorphous domain surrounding the nacre tablets (15-20 nm thick) composed of organic matter rich in calcium and deficient in carbonate [35], which support the presence of calcium-binding acidic proteins.

**Figure 5.**
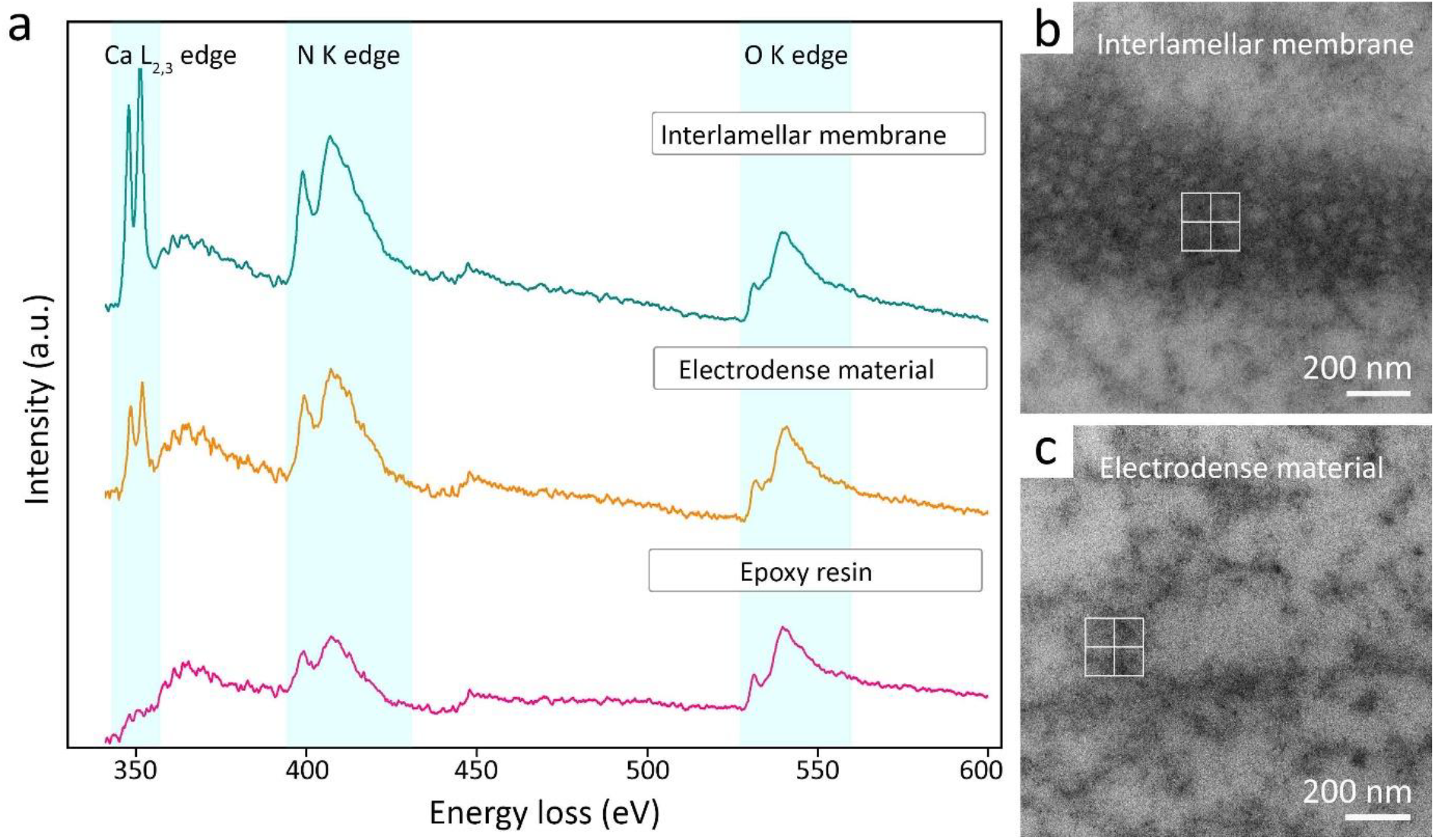
Calcium is bound to protein complexes. a) Spectra from the two organic structures in the mineralization chamber (interlamellar membranes and electrodense material) compared against the epoxy resin, showing an increasingly higher content of calcium and nitrogen. b-c) TEM images of the sampled areas.

**Figure 6.**
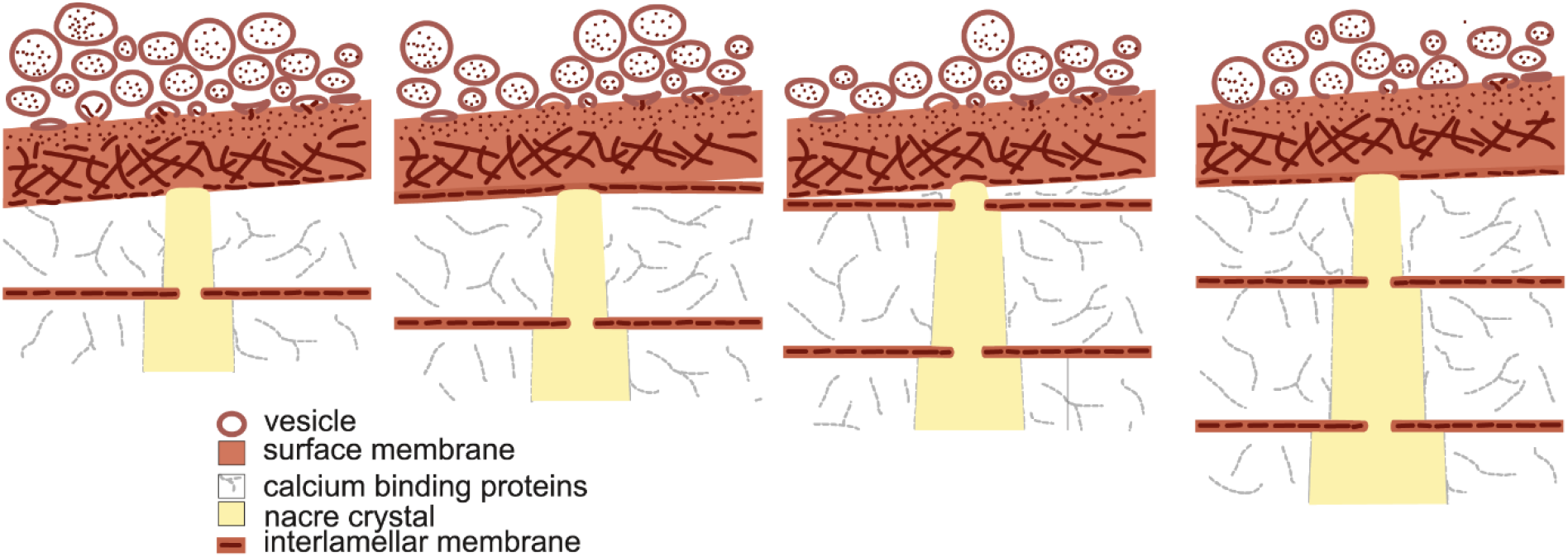
Polymerization of surface membrane components and formation of interlamellar membranes. Model representing the formation of the surface membrane by the addition of vesicles and their cargo, and the structural changes through its thickness, from a disordered structure to an intermediate fibrillar structure. Once the final porous structure is reached, the interlamellar membranes separate from the surface membrane as the aragonite platelet grows in heigh.

Our data demonstrated that the insoluble fraction that forms the surface membrane and interlamellar membranes bound a significant amount of calcium in the form of calcium-protein complexes prior to mineralization. The preparation techniques used in this study did not allow for the detection of amorphous calcium carbonate (ACC), and in fact, we could not completely exclude the loss of soluble calcium. However, it is noteworthy that hydrated ACC was found distributed along the interlamellar membranes [10] and at the border of the growing tablets [35-37]. Therefore, our results indicate the presence of a transport mechanism based on calcium-binding proteins. It remains to be known whether these or similar proteins also bind ACC or whether organic calcium reacts with carbonate ions in a subsequent step.

## Acknowledgement

EMS is supported by the Knowledge Generation Project PID2022-141993NA-I00 funded by MICIU/AEI/10.13039/501100011033 and FEDER (EU) and the Research Program Juan de la Cierva Incorporación (IJC2020-043639-I) funded by MCIN/AEI/10.13039/501100011033 and EU NextGenerationEU/PRTR. A.G.C. acknowledges funding by projects PID2020116660GB-I00 (Spanish Ministry of Science and Innovation: MCIN/AEI/10.13039/501100011033/) the Unidad Científica de Excelencia UCE-PP2016-05 (University of Granada) and the Research Group RNM363 (Junta de Andalucía).

