## Supplementary material for "Organic macromolecules transport a significant proportion of the calcium precursor for nacre formation"

[5] Instituto Andaluz de Ciencias de la Tierra, CSIC–Universidad de Granada, Armilla, Spain

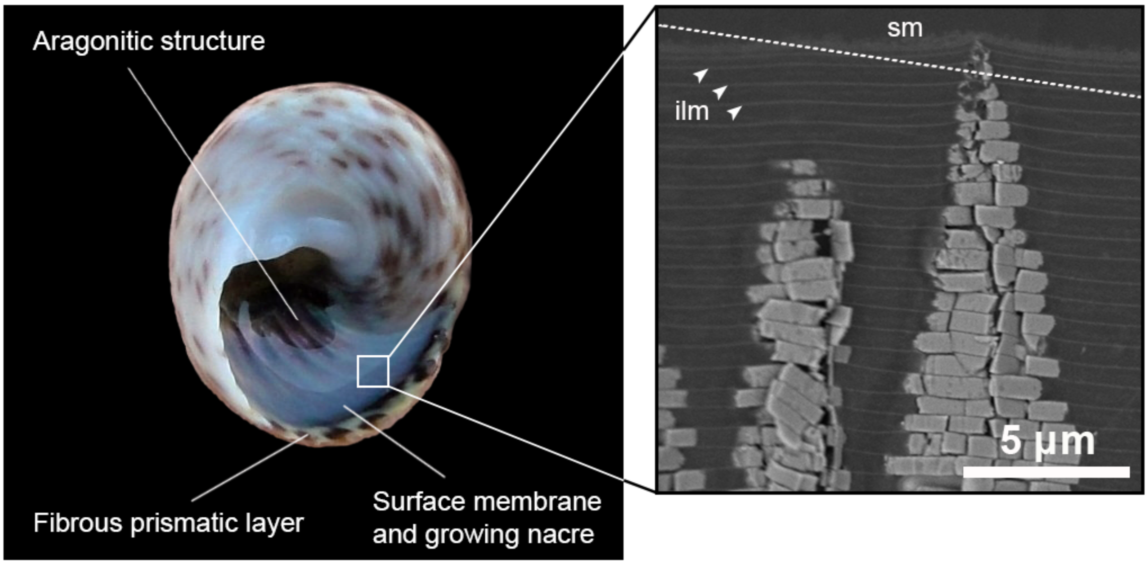


**Figure S1.** A) Left, specimen of *Phorcus turbinatus* indicating the location of the surface membrane and the growing nacre (sampled area) in relation to the other microstructures present at the shell aperture. B) SEM image of an ultramicrotome slice perpendicular to the shell surface, where it is possible to distinguish the surface membrane (sm), the interlamellar membranes (ilm) and two towers of growing nacre. The dotted line indicates the approximate angle of the ultramicrotome cuts carried out in this study.

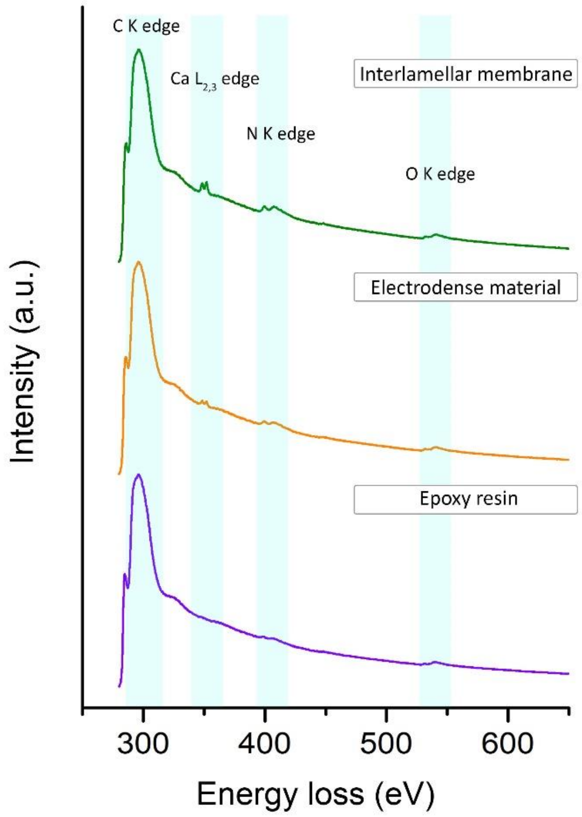


**Figure S2. EEL spectra from the three main regions.** Each spectrum corresponds to the sum of two spectra after background subtraction, no smoothing was applied. Intensity is normalized and expressed in arbitrary units (a.u.) to facilitate comparison.

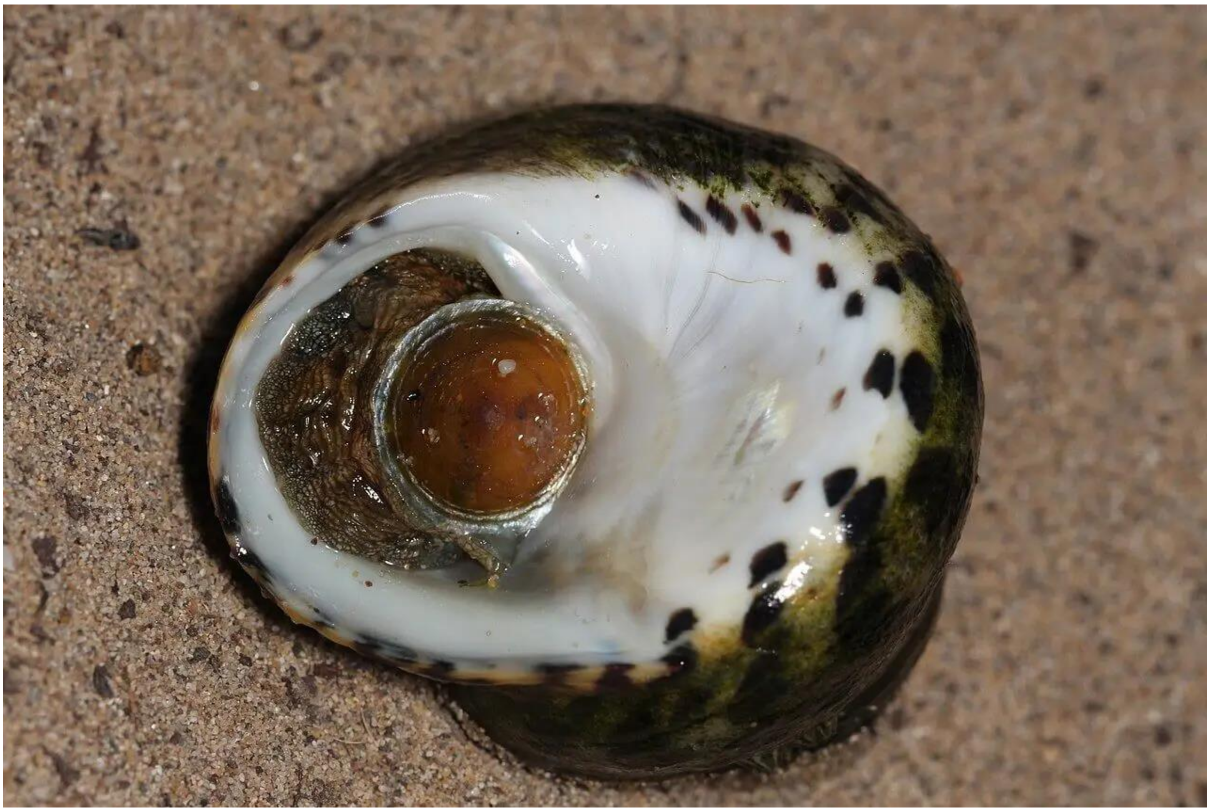

**Figure S3.** Image showing the retraction capacity of the gastropod body, closing the entrance opening with the operculum and leaving the nacre growth area and the surface membrane in contact with seawater.
